# A novel event improves memory retrieval and divergent thinking in a naturalistic school environment

**DOI:** 10.64898/2026.03.05.709820

**Authors:** D. Ramirez Butavand, A. Barbuzza, P. Bekinschtein, F. Ballarini

## Abstract

Stored memories are useless unless they are available for retrieval. Thus, investigating different ways to modulate retrieval is crucial. Novelty has been extensively studied as a modulator of memory. In this study, we investigated whether exposure to a novel event, an innovative neuroscience lesson, can enhance memory retrieval and divergent thinking in high school students. Across three experiments, we assessed the timing and mechanisms underlying these effects. In experiment 1, we found that memory retrieval was enhanced when the novel lesson occurred immediately before a memory test, but not when it was presented one hour earlier. In experiment 2, we found that the same immediate novelty exposure improved divergent thinking performance. Finally, in experiment 3, we explored potential shared mechanisms using a competition protocol and revealed that novelty improved divergent thinking regardless of its timing relative to memory retrieval. However, memory retrieval benefited only when tested immediately before the divergent thinking task. These results suggest that novelty boosts both memory retrieval and divergent thinking, but through partially distinct mechanisms. Our findings demonstrate that a simple, real-world classroom intervention can effectively enhance key cognitive functions in students.

**Significance Statement:** Stored memories are only valuable if they can be retrieved, and memory retrieval plays a key role in creative thinking. Here, we tested whether a simple, novel event, a neuroscience lesson, could enhance memory retrieval and creative thinking in a real-world classroom setting. We found that novelty improved both memory retrieval and divergent thinking, an aspect of creative thinking, when presented immediately before the task. Finally, we revealed a non-reciprocal competition effect between memory retrieval and divergent thinking. These findings highlight a practical, low-cost intervention to boost key cognitive functions in students, demonstrating that brief, well-timed novel experiences can support both learning and creative thinking in educational environments.

## Introduction

Learning is fundamental to both survival and social progress. However, stored memories are only useful if they can be accessed at the appropriate time. Memory retrieval involves the access, selection, and reactivation of stored internal representations (1). While the mechanisms underlying learning and memory storage have been extensively studied, the processes that govern memory retrieval and possible strategies to enhance it remain less understood. Various internal states, such as arousal, stress, motivation, or reward, are known to influence memory formation (2-6). Furthermore, over the last decades, evidence from both animal and human studies has shown that exposure to novel events close to learning can also modulate memory (7). Specifically, novelty has been found to enhance memory consolidation when presented within a critical time window around learning (8-10). However, the potential for novelty to modulate memory retrieval has been less studied—especially in humans—with only a few reports available from rodent models. Understanding this effect in real-world settings, such as schools, could offer a valuable and practical tool for enhancing learning outcomes.

Memory retrieval is essential for several cognitive functions, including recognition, decision-making, and creative thinking (11-12). Creativity is often defined as a cognitive process that relies on the rapid combination and recombination of internal mental representations to generate novel ideas and ways of thinking (13). It is now widely accepted that creativity is a universal human trait and a hallmark of cognitive flexibility (14), involving the production of ideas that are both original (i.e., rare and unexpected) and appropriate (15-16). Through creativity, humans have transformed their environment and achieved significant accomplishments, developed innovative solutions to complex problems, and advanced intellectually and culturally (17-18).

One measurable component of creativity is divergent thinking (19). This process requires mental flexibility, enabling the generation of multiple possible solutions in contexts where selection criteria are vague and more than one answer may be correct (20-21). For this process, it is essential to access existing mental representations stored in memory (13). Although memory retrieval and divergent thinking were traditionally studied independently, research over the past decade has increasingly linked the two. Divergent thinking has been associated with hippocampal function, as individuals with bilateral hippocampal lesions show poorer performance on divergent thinking tasks compared with healthy controls (23). In addition, enhancing the recall of specific details from recent experiences through an episodic-specific induction has been shown to improve divergent thinking performance, suggesting that this form of creativity relies on episodic memory processes (24).

To further investigate the relationship between memory retrieval and creative thinking, particularly within an educational context, we aimed at determining whether exposure to novelty could enhance both processes in high school students. Specifically, we examined whether a novel event improved memory retrieval and divergent thinking, and whether these effects relied on shared underlying mechanisms. Understanding how novelty influences these cognitive functions, and how they interact, could provide valuable insights for developing simple, effective educational strategies to boost both memory performance and creativity in the classroom.

## Materials and Methods

### Participants

A total of 414 students (125 females), aged 12 - 16 years (mean 14.1 ± 1.2 years), participated in the study. Participants were recruited from seven different high schools in Buenos Aires, and were naïve to the experimental procedures. This study was approved by the Ethical Committee of the *Instituto de Tisioneumonología “Prof. Dr. Raúl Vaccarezza”* of the University of Buenos Aires. Approval was also obtained from the head of each participating institution. Written informed consent was provided by the students’ parents or legal guardians, and participation was voluntary; students could withdraw from the study at any time without consequences.

### Procedure

#### Experiment 1

To evaluate the effect of novelty on memory retrieval (Figure **1a**), participants first completed a visual memory encoding task. Two days later, memory retrieval was tested. The novelty groups received a novel lesson immediately before (-0 h) or one hour prior (-1 h) to retrieval, while the control group attended their regular school lessons during this period.

**Figure 1:**
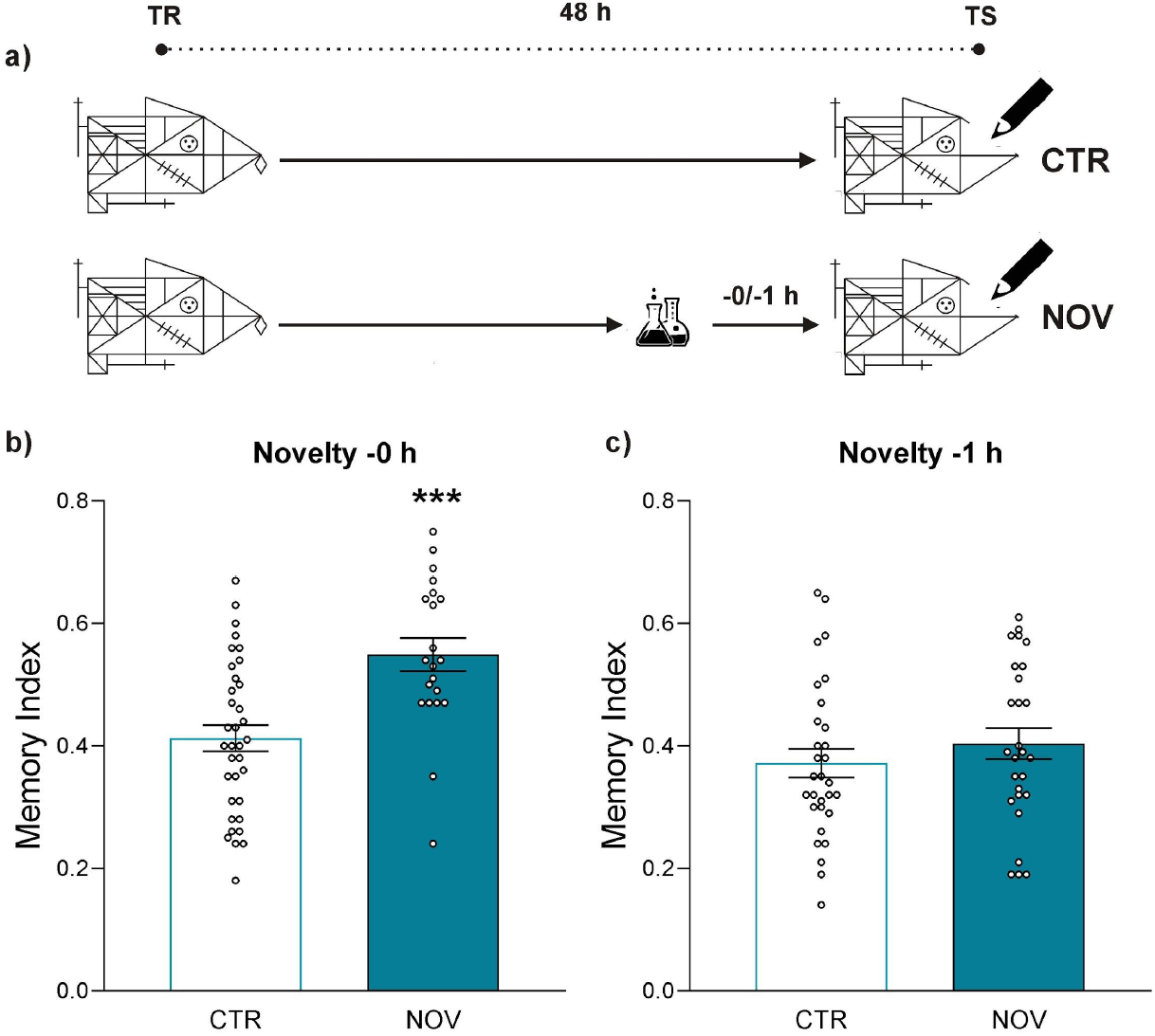
A novel event improves memory retention only when it is given immediately before test. **a)** Schematic representation of the experimental protocol used. The novelty was given immediately before (-0 h) or one hour before (-1 h) the test (TS) of the memory formed during the training (TR) session two days earlier. Rey-Osterrieth’s figure memory indices are shown as the mean ± SEM for both the control (CTR) and novelty (NOV) groups. **b)** The novel lesson was given immediately before the memory test. The number of participants in the CTR group was 35 and in the NOV group 21. Student’s *t*-test CTR vs NOV: *t*(54) = 3.96, *** *p* < 0.001. **c)** The novel lesson was given one hour before the memory test. The number of participants in each group was 30 and 27, respectively. Student’s *t*-test CTR vs NOV: *t*(55) = 0.93, *p* = 0.356.

#### Experiment 2

To assess the impact of novelty on divergent thinking (Figure **3a**), the novelty group attended a novel lesson immediately before completing a divergent thinking task. The control group completed the same task following their usual school activities.

#### Experiment 3

This experiment examined the effects of novelty on memory retrieval and divergent thinking when both tasks were performed in sequence. All participants completed the memory encoding task on day one. Two days later, groups followed different protocols: one group performed memory retrieval followed immediately by the divergent thinking task; a second group received a novel lesson before performing these tasks in the same order. A third group completed the divergent thinking task first, followed by memory retrieval; the fourth group followed this same order but received the novel lesson before starting the first task (Figure **4a**).

For additional control, pairs of groups performed a non-creative task before memory retrieval (Figure **5a**). The novelty group received the novel lesson immediately before the non-creative task, while the control group did not.

### Intervention

#### Novel lesson

To qualify as a novel event, the activity met the following criteria: (1) students were unexpectedly taken from their regular classroom to a different, unfamiliar location within the school; (2) this location was not typically used for lessons; (3) the lesson was delivered by a trained member of the research team unknown to the students; (4) it was a brief (20-minute) interactive session, never previously experienced by the students, featuring age-appropriate novel content; and (5) students were encouraged to actively participate and maintain attention throughout. After the activity, students returned to their usual classroom.

The novel lesson focused on *Attentional Blindness* and included simple experiments designed to engage students fully. They participated in games demonstrating how people often overestimate their attentional capacity. Two main phenomena were explored: **change blindness**, where viewers fail to notice substantial changes in images (using materials from 23); and **inattentional blindness**, where obvious elements are overlooked due to focused attention on a task (using videos from 24). The lesson concluded with a brief overview of basic biological and neuroscientific principles related to the experiments.

Notably, such an interactive attentional activity, without subsequent testing on the content, is uncommon in high school curricula. This novel intervention has been successfully used in prior studies on the effects of novelty on memory consolidation (8-9).

### Experimental Tasks

#### Visual Memory Task

To assess memory performance, we used a standardized complex figure composed of 18 geometric elements arranged into a unified design (6-8,25).

The protocol spanned two days. On the training (TR) day, each student received a printed copy of the figure and was instructed to reproduce it by hand on a blank sheet placed beneath the model, within a 5-minute time limit, sufficient to copy the entire figure. Long-term memory (LTM) was evaluated two days later, on the testing (TS) day, when students were asked to draw the figure from memory on a blank within a 5-minute interval.

Drawings were scored following the Rey-Osterrieth system, which assesses each of the 18 elements for location, accuracy, and organization (26). Briefly, **0 score (null)**, when the element is absent; **0.5 score (low)**, when the element is present but in the wrong location and with errors; **1 score (mid)**, when the element is correctly drawn but in the wrong location, or partially correct in the correct location; and **2 score (high)**, when the element is correctly drawn and correctly located.

For each student, we calculated a *Memory Index* as the ratio between their score on the testing day and their score on the training day. Also, we quantified the proportion of items in each category (null, low, mid, and high) relative to the total number of elements.

#### Divergent Thinking Task: Alternate Uses Task (AUT)

To assess divergent thinking, we used the Alternate Uses Task (AUT) (27) in which participants were asked to list as many possible uses for a paper clip as they could within 5 minutes. Before starting, they were given a simple example to ensure understanding.

Performance was evaluated following a standardized scoring method (22) across two dimensions: fluency and originality. Fluency was calculated as the total number of relevant, non-redundant uses generated (1 point per valid use). Originality was calculated as the total score based on how uncommon each valid use was within the participant’s experimental group (< 5% occurrence = 2 points; 5–10% = 1 point; > 10% = 0 points). For each participant, scores from fluency and originality were summed to obtain the Alternative Uses Index.

#### Control Task: non-creative version

In this task, participants received a list of potential uses for a paperclip (e.g., opening a padlock -possible; taking a picture -not possible). For each item, participants marked whether the proposed use was **possible, not possible**, or if they **did not know**. The list included space for these responses, and participants had 5 minutes to complete it. This task was not scored, as its purpose was to serve as a non-creative control, maintaining a similar format and timing to the divergent thinking task but without requiring the generation of novel ideas.

### Statistical Analysis

In the results of experiments where the effect of a novel lesson was evaluated in a single task, the groups were compared using Student’s *t*-test. When competition between two tasks was assessed or when performance was differentiated by gender, we used a 2-way ANOVA. In cases where there were significant interactions, a post hoc analysis of multiple Sidak comparisons was performed. Participants who were two standard deviations away from the mean on each task were excluded. Statistical significance was determined using an alpha level of **α** = 0.05. The results were analyzed with the software GraphPad Prism®.

## Results

### Experiment 1

In Experiment 1, we tested whether a brief novel lesson could enhance long-term visual memory retrieval in high school students. Participants first copied a complex figure during a training session (TR), then reproduced it from memory 2 days later during a memory retrieval test (TS). Before the retrieval test, novelty groups attended a 20-minute novel lesson either immediately before (-0 h) or one hour before (-1 h) the task, while control groups followed their regular school lesson (Figure **1a**). Memory performance was quantified using the Memory Index (see Methods).

We found that the group that received the novel lesson immediately before the memory test showed significantly higher memory performance than the control group (Figure **1b**, *t*(54) = 3.96, *** *p* < 0.001, 95% CI [0.067, 0.205]), Cohen’s d = 1.09) whereas the -1h novelty group performed similarly to the control group (Figure **1c**, *t*(55) = 0.93, *p* = 0.356, 95% CI [-0.037, 0.100]).

Finally, we examined the proportion of scores assigned to each element of the figure, following the approach of previous studies (6). For each participant, the number of items in each scoring category (Null, Low, Mid, High) was divided by the total number of elements (18).

In the immediate-novelty condition (Figure **2a**), the higher overall performance in the NOV group was mainly explained by fewer undrawn items (Null: t(208) = 5.21, *** p < 0.001) and more correctly drawn and correctly placed items (High: *t*(208) = 6.45, *p* < 0.001) compared with controls. No differences were found for partially correct items (Low: *t*(208) = 1,75, *p* = 0.307; Mid: *t*(208) = 0.79, *p* = 0.899).

**Figure 2:**
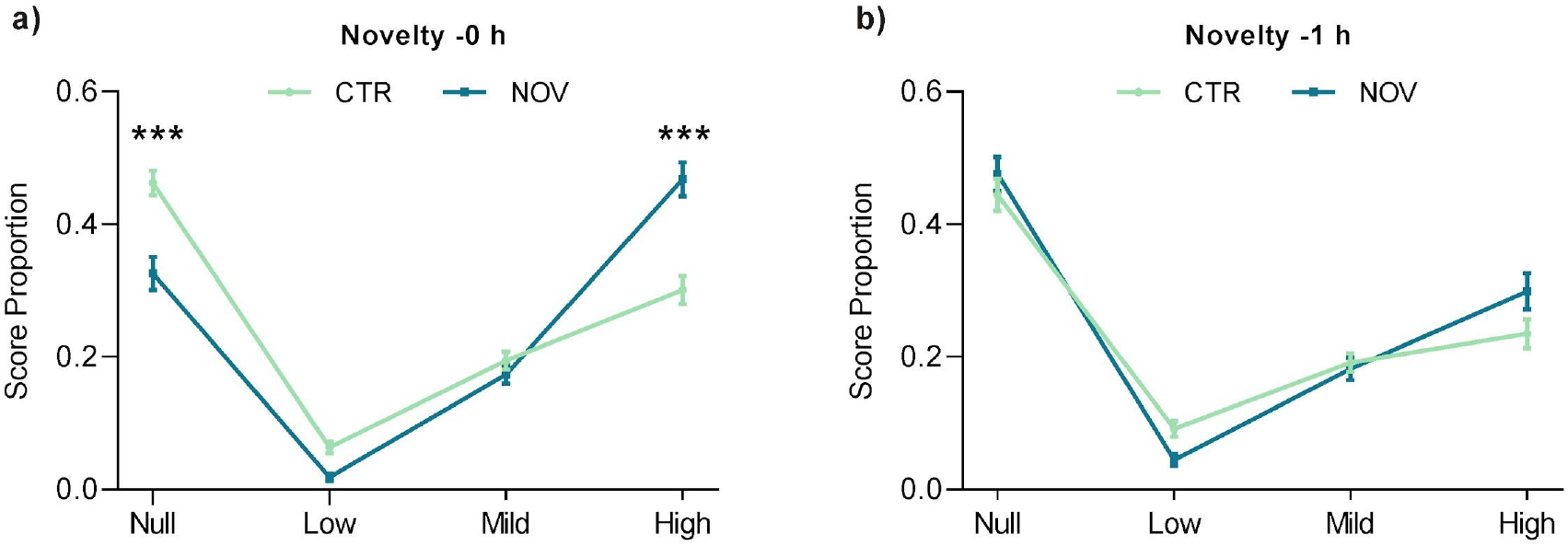
Score proportion for the graphic memory. The score proportion obtained is shown as the mean ±SEM for the control and novelty groups in each possible score: Null, Low, Mid, and High. **a)** We compared the profile of the control (green) and the novelty -0 h (blue), obtaining significant differences in the Null score (*t*(208) = 5.21, *** *p* < 0.001) and the High (*t*(208) = 6.45, *** *p* < 0.001). In the Low and Mid scores, no significant differences were observed between the groups *p* > 0.05. **b)** We compared the profile of control (green) and novelty -1 h (blue) and found no significant differences between groups in any of the possible scores (*p* > 0.05 in all cases).

In the –1 h novelty condition (Figure **2b**), which did not improve memory performance (Figure **1c**), both groups showed similar score profiles (Null: *t*(212) = 1.13, *p* = 0.703; Low: *t*(212) = 1.67, *p* = 0.336; Mid: *t*(212) = 0.05, *p* > 0.999; High: *t*(212) = 2.30, *p* = 0.086).

### Experiment 2

In Experiment 2, we evaluated whether the same novel lesson could enhance divergent thinking, measured with the Alternative Uses Task (AUT). Participants listed as many uses for a paperclip as they could in 5 minutes. Performance was scored in two dimensions, fluency and originality, and summed to obtain the “Alternative Use Index”. The control group (CTR) completed the AUT without prior intervention, while the novelty group (NOV) received the novel lesson immediately beforehand (Figure **3a**).

**Figure 3:**
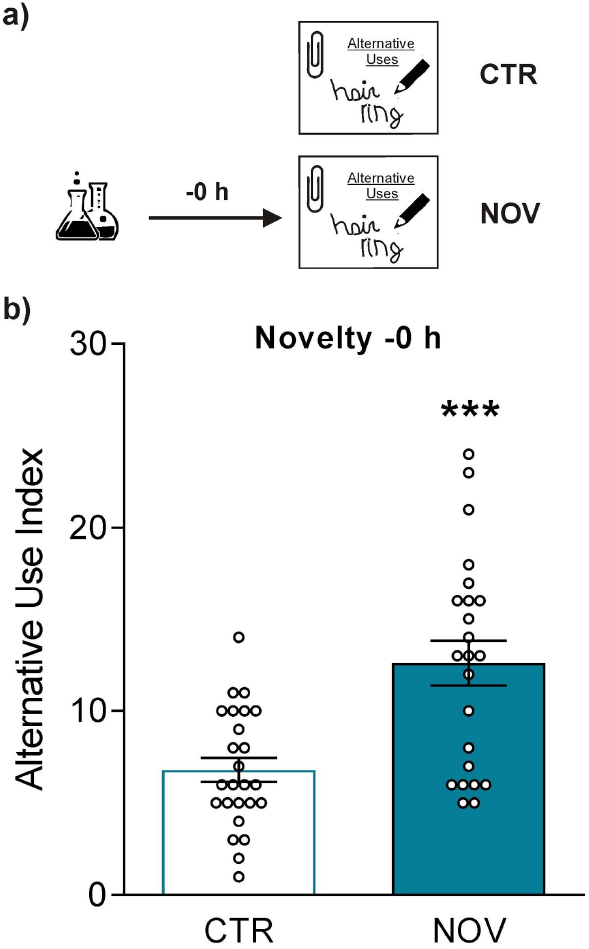
A novel lesson immediately before the alternative uses task improves performance. **a)** Schematic representation of the experimental protocol used. The novel lesson was given immediately before the alternative uses task. **b)** Total Clip, calculated by summing fluency and originality for each course, is shown as the mean ± SEM. The number of participants in each group was 25 and 23, respectively. Student’s *t*-test: *t*(46) = 4.32, CTR vs. NOV, *** *p* < 0.001.

We found that the NOV group achieved a significantly higher Alternative Use Index than the CTR group (*t*-test comparison (*t*(46) = 4.32, *** *p* < 0.001, 95% CI [3.100, 8.517], Cohen’s d = 1.25).

### Experiment 3

Given that a novel event enhanced both memory retrieval and divergent thinking when tested separately, in Experiment 3, we examined whether these effects relied on shared resources or occurred independently. Each group completed both tasks consecutively: one group retrieved the figure learned two days earlier and then performed the AUT, another group completed the tasks in the reverse order, and two additional groups followed the same protocols but received the novel lesson immediately before the first task (Figure **4a**).

**Figure 4:**
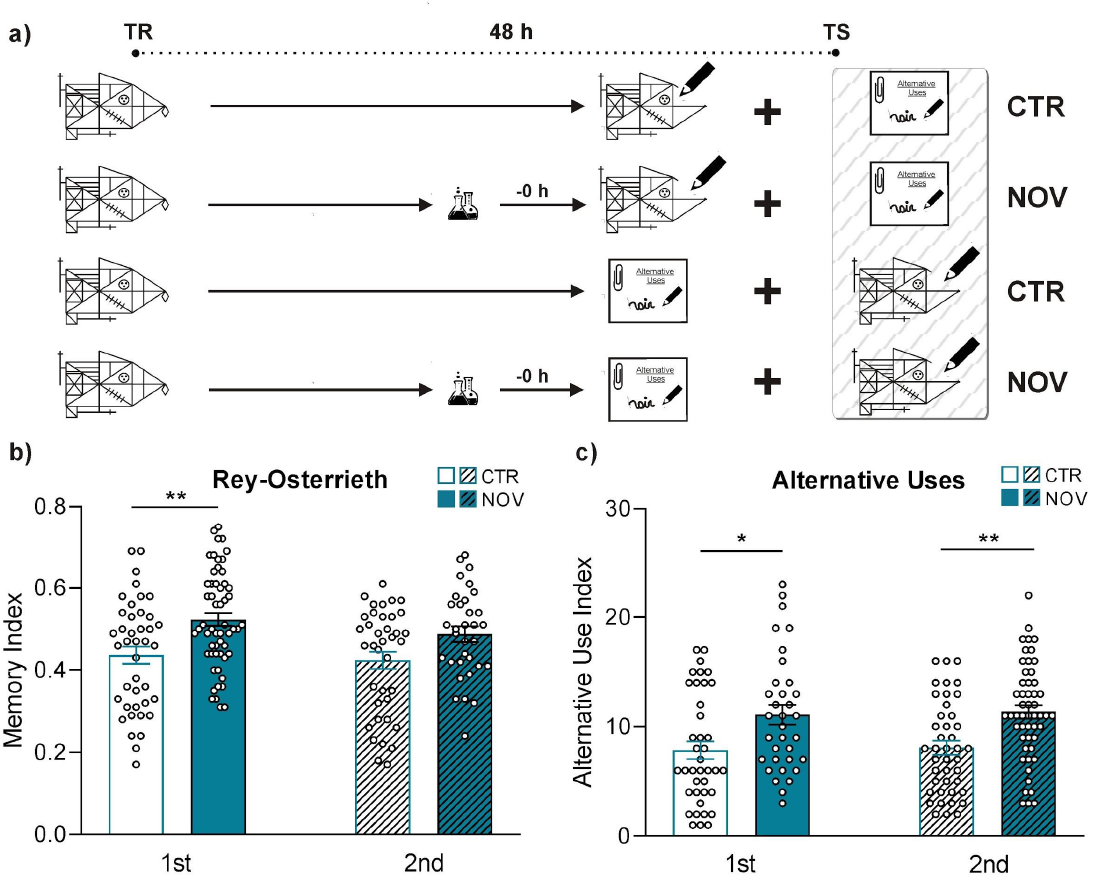
Retrieval - AUT interaction. **a)** Schematic of the experimental protocol used to study the competition between memory retrieval and divergent thinking processes. **b)** Rey-Osterrieth’s figure memory indices for each group are shown as the mean ± SEM. The 2-way ANOVA revealed a significant main factor (Novelty), (Novelty Factor: F(1,164) = 15.29, *** *p* = 0.001, ηp2=0.085), and the post hoc analysis reflected a significant difference between the control and novelty groups only if the memory retrieval test was performed first (t(164) = 3.46, ** *p* = 0.004). The number of participants in each group was: 41, 56, 38, 33. **c)** The Total Clip score of the AUT was calculated by summing the fluency and originality and is shown as the mean ± SEM. The 2-way ANOVA analysis reflected a significant difference in its main factor (novelty), i.e., irrespective of the order in which the AUT was performed, the novelty group showed better performances (Novelty factor: F(1,166) = 22.80, *** *p* < 0.001, ηp2=0.121). Post hoc analysis revealed that performance in AUT improved regardless of task order: t(166) = 3.44, ** *p* = 0.004 when the AUT was performed before the memory task, and t(166) = 3.32, ** *p* = 0.006 when it was performed after the memory task. The number of participants in each group was: 39, 34, 42, 52. The solid bars correspond to the task that was performed first, and the striped bars to the task that was performed second.

For the memory task, a 2-way ANOVA revealed a significant main effect of novelty (*F*(1,164) = 15.29, *** *p* < 0.001, ηp2=0.085), but no significant main effect of order (*F*(1,164) = 1.50, *p* = 0.221) and no significant interaction (F(1, 164) = 0.3746, p = 0.541). A Post-hoc analysis showed that the NOV group outperformed the CTR group only when the memory task was performed first (Figure **4b**, *t*(164) = 3.46, ** *p* = 0.004), replicating the effect seen in Experiment 1 (Figure **1b**). When the memory test followed the AUT, no difference was found between groups (*t*(164) = 2.18, *p* = 0.133).

For the AUT, a 2-way ANOVA revealed a significant main effect of novelty (*F*(1,166) = 22.80, *** *p* < 0.001, ηp2=0.121), but no significant main effect of order (*F*(1,166) = 0.51, *p* = 0.475) and no significant interaction (*F*(1,166) = 0.12, *p* = 0.733). However, the post hoc analysis NOV vs CTR revealed that performance improved regardless of task order: (Figure **4c**, 1st: t(166) = 3.44, ** *p* = 0.004; 2nd:t(166) = 3.32, ** *p* = 0.006).

To explore why the novel lesson did not boost memory retrieval when it followed the AUT, we conducted a control experiment in which the AUT was replaced with a similar but non-creative task. Participants first completed this task, marking a list of paper clip uses as “possible”, “not possible”, or “don’t know”, and then tested for memory of the Rey Ostherrieth’s figure learned two days earlier. The NOV group received a novel lesson immediately before the non-creative task (Figure **5a**).

**Figure 5:**
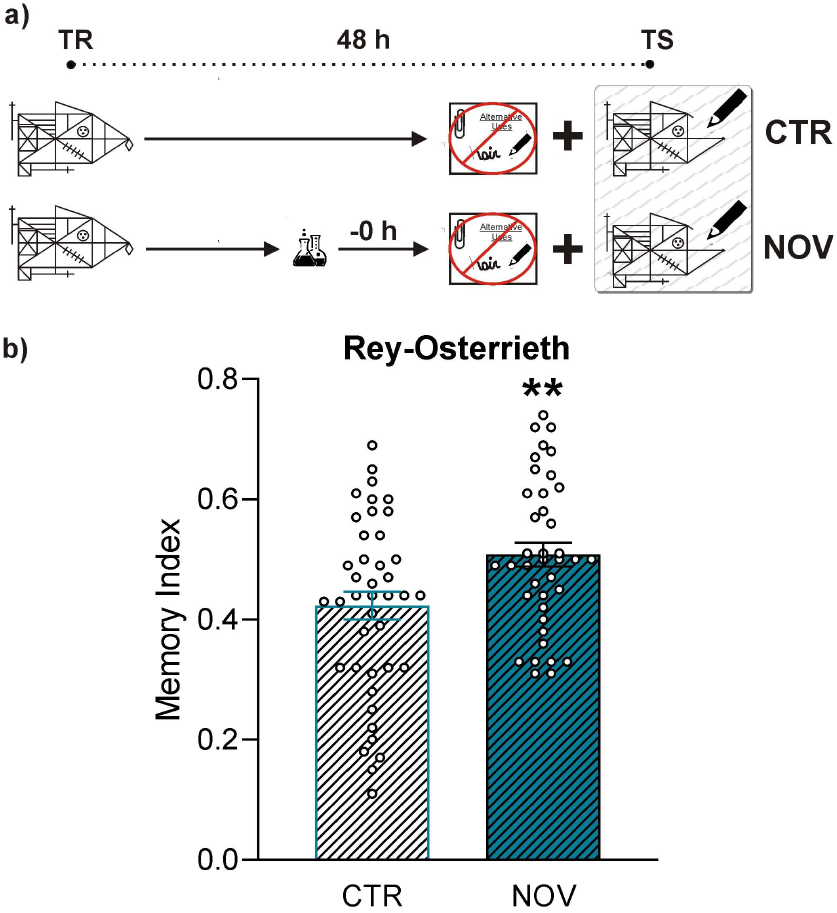
There is no competition between the non-creative task and memory retrieval. **a)** Experimental protocol conducted to evaluate the interaction between the non-creative version of the alternative uses task and memory recall. **b)** Rey-Osterrieth’s figure memory indices for each group are shown as the mean ± SEM. The number of participants in each group was: 41, 39. Student’s *t*-test CTR vs NOV: t(78) = 2.75, ** *p* = 0.007.

When the memory retrieval test followed the non-creative task, performance was significantly higher in the NOV group than in the CTR group (Figure **5b**, *t*(78) = 2.75, ** *p* = 0.007; 95% CI [0.023, 0.146], Cohen’s d = 0.62).

## Discussion

In this study, we investigated how a novel event influences memory retrieval and divergent thinking in an ecological educational context. We found that a novel event enhanced memory retrieval only when it was presented immediately before the test, but not when delivered one hour earlier. Novelty also improved performance on the divergent thinking task (AUT) when presented immediately before the task. To determine whether these two processes interact, we analyzed their performance when carried out consecutively. The novel event enhanced divergent thinking regardless of task order, whereas its beneficial effect on memory retrieval appeared only when retrieval occurred before the AUT.

Previous results have shown that a novel event presented around learning can improve long-term memory of an unrelated graphical memory in primary and high school students (8-9). Here, we assessed whether novelty can also modulate memory retrieval. Evidence for such an effect has been scarce, with one study with rodents (28) reporting that exploration of a novel open field enhanced retrieval of a fear memory when administered 0 to 2 hours before testing. Our results indicate that novelty also facilitates episodic memory retrieval in humans, but within a narrower temporal window: enhancement was observed only when the novel event occurred immediately before testing. This difference is not unexpected given that the type of memory, the nature of the novel experience, and the testing context differed substantially between species.

For each student, we quantified the number of items receiving null, low, medium, or high scores across experimental conditions. When the novel event enhanced memory retrieval (novelty -0 h), students omitted fewer elements and produced more items drawn accurately and in the correct position compared with the control group. However, when the novel event had no effect (novelty -1 h), the score distributions were indistinguishable from those of the control group. This pattern indicates that novelty primarily improved the precise recall of both the composition and location of the figure’s elements, closely resembling the score profiles observed in a previous study in which a school exam improved consolidation of Rey-Osterrieth’s figure (6).

The novel event also improved the performance on the divergent thinking task. Creativity is likely one of the key aspects to develop in education, as it depends on both the availability of relevant information and its flexible recombination to generate novel ideas (13-16). Understanding how memory retrieval (information availability) and divergent thinking (idea generation) interact in the classroom is therefore highly relevant for improving learning outcomes. Several studies conducted during the 1960s reported positive associations between creativity and academic performance (29-30), and more recent research in elementary school (31) and college students (32) has confirmed that creativity predicts academic performance. Therefore, identifying simple strategies, such as brief novel events, that reliably enhance creativity may have practical value in education.

Our interaction experiments revealed an asymmetric pattern: the novel event improved divergent thinking, whether it occurred before or after the memory test, whereas memory retrieval improved only when it preceded the AUT. Notably, when the novel event occurred before the AUT and memory retrieval followed, there was a trend toward better memory performance, although this difference was not statistically significant. This pattern suggests that the AUT may partially consume the cognitive or neural resources engaged by the novel event, reducing the opportunity for novelty to benefit subsequent memory retrieval.

A plausible explanation is that the divergent thinking task draws on the same novelty-sensitive resources needed to enhance memory retrieval. Support for this interpretation comes from the control experiment in which the AUT was replaced with a similar non-creative task: under these conditions, the novel event again boosted memory retrieval, indicating that the creative component of the AUT was responsible for the competition. This idea is consistent with neuroimaging research showing that both episodic retrieval and creative idea generation recruit overlapping brain networks, including the hippocampus, medial temporal lobe structures, lateral and medial parietal regions, and medial prefrontal cortex (33-34). Increased activation in these regions has also been observed when participants generate creative ideas (35-36).

However, divergent thinking improved after the novel event regardless of whether it was performed before or after the memory test. If both processes rely on overlapping novelty-sensitive resources, this asymmetry suggests that retrieval of the Rey–Osterrieth figure may consume fewer of these resources than the AUT. Thus, even when memory retrieval is completed first, sufficient resources may remain for the AUT to benefit from the novel event. In contrast, when the AUT is performed first, its higher cognitive and associative demands may tax these shared resources more heavily, leaving little available to enhance subsequent memory retrieval. Nevertheless, because these functional demands have not been directly compared in a way that maps onto the present behavioral effects, further experiments will be required to determine whether differential resource consumption truly underlies the observed asymmetry.

Although the mechanisms underlying novelty-induced enhancement of memory retrieval were not directly assessed in the present study, neuromodulatory pathways, particularly dopaminergic and noradrenergic systems, represent plausible candidates. Pharmacological manipulations of dopaminergic transmission have been shown to modulate episodic memory retrieval. Acute administration of a dopaminergic D2 receptor antagonist (haloperidol) enhanced recall performance and increased activity in retrieval-related regions, including the striatum, prefrontal cortex, hippocampus, SN/VTA, and anterior cingulate cortex (37). Conversely, reduced dopaminergic transmission impairs item-recognition retrieval in humans (38). These findings suggest that heightened dopaminergic signaling facilitates access to stored information. Because novel events reliably trigger dopaminergic and noradrenergic activation (39) and because retrieval itself depends on noradrenergic mechanisms (40), a novelty-evoked increase in these neuromodulators could jointly support the boost in long-term memory retrieval observed for the Rey–Osterrieth figure.

A similar neuromodulatory mechanism could account for the observed improvement in divergent thinking. Chermahini and Hommel (41) reported an inverted-U relationship between spontaneous eye-blink rate (an indirect proxy of dopamine levels) and divergent thinking performance. This suggests that optimal, intermediate dopamine levels promote flexible recombination of stored information. If the novel event elevates dopamine into this optimal range, it could enhance both memory retrieval and divergent thinking, providing a unified physiological pathway through which novelty supports cognitive performance. Further work will be required to directly test this hypothesis.

We live in a complex and constantly evolving world that demands continual innovation and flexible problem-solving. Among the key competencies for navigating the 21st century, creativity stands out as a core educational skill (13-16). At the same time, memory constitutes a cornerstone of learning, as new knowledge can only be integrated through its connection with prior information and past experiences (42-43). In this study, we show that both divergent thinking and memory retrieval can be enhanced in an ecological school setting through a simple, low-cost intervention: the introduction of a brief novel event. Novelty is easy to implement, adaptable to different curricular contents, and does not require additional resources (8-9). Although frequent use may reduce its effectiveness as novelty becomes predictable, its flexibility allows targeted application. For instance, teachers may strategically introduce novelty before specific lessons to strengthen the retrieval of particular content, or use it before activities in which divergent thinking is especially desirable. Overall, these findings provide initial evidence that incorporating well-timed novel experiences into classroom practice may offer a practical, evidence-based approach for enhancing both learning and creative thinking from early stages of education.

## Data Availability

Upon publication, all data will be made publicly available at the PsychArchives of the Leibniz Institute of Psychology (ZPID) and will be accessible at a persistent DOI. We can provide preview-access to the reviewers. This study was not pre-registered.

## Acknowledgments

We thank the schools that participated in this research (San Carlos Borromeo, Modelo Lomas, Escuela Superior de Comercio Carlos Pellegrini, San Agustín, and Instituto Mario Madeddu), and we are especially grateful to the iTINERE educational network and its General Director, Darío Álvarez Klar.

## Conflict of interest

The authors declare that the research was conducted in the absence of any commercial or financial relationships that could be construed as a potential conflict of interest.

## Notes

### Competing Interest Statement

The authors have declared no competing interest.

